# Latent Structure of Risk Perception

**DOI:** 10.1101/389890

**Authors:** Elisa Pabon, James MacKillop, Abraham A. Palmer, Harriet de Wit

## Abstract

Risk-taking behavior affects many aspects of life, including maladaptive behaviors such as illicit substance use, unsafe driving, and risky sexual behavior. Risk-taking has been measured using both self-report measures and behavioral tasks designed for the purpose, but there is little consensus in the associations among measures and our understanding of the latent constructs underlying different forms of risk is limited. In the present study we examined the construct of risk using data from over 1000 young adults who completed measures of risk-taking, including self-reports of perception of risk, propensity to engage in risky behaviors and performance on behavioral tasks designed to measure risk. To examine the latent structure of risk preferences, we conducted a principal component analysis (PCA). The PCA revealed a latent structure of three distinct components of risk-taking behavior: “*Lifestyle Risk Sensitivit*y”, “*Financial Risk Sensitivity”*, and “*Behavioral Risk Sensitivity”,* which consisted only of the Balloon Analogue Risk Task (BART; Lejuez et al., 2002). As expected, risk-taking and perception of risk differed in men and women. Yet, the PCA components were similar in men and women. Future work utilizing additional measures of risk-taking behavior in more heterogeneous samples will help to identify the true biobehavioral constructs underlying these behaviors.

## 1. Introduction

Many of life’s decisions focus on opportunities to gain some reward but with the possibility of a potential loss or other possible harm (Leigh et al. 1999). Such decisions are commonly referred to as ‘risk-taking behaviors’, typically involving voluntary engagement in reward-seeking activities that are probabilistically linked to monetary, social or interpersonal loss (Bechara, 2003). Until recently, risk-taking was considered to be a relatively discrete personality trait, and individuals would be categorized as risk-taking or risk averse (Eysenck & Eysenck, 1977; Bromiley & Curley, 1992; Lejuez et al., 2002). For example, a recent genetic study, which included over one million individuals, utilized a single item to assess risk tolerance, in essence “Would you describe yourself as someone who takes risks? Yes / No” (Linnér et al., 2018) However, further empirical evidence suggests that risk-taking may not be a unitary construct, but instead multidimensional, and vary across domains, including financial, ethical and social Duijvenvoorde et al. 2015, Zuckerman and Kuhlman, 2000; Blais and Weber, 2001, Horvath and Zuckerman, 1993).

In a recent study, Frey et al (2017) summarize the challenges in studying ‘risk’ as a construct, including questions of whether the tendency to take risks is unitary or multidimensional, or stable or changeable. They note that risk is defined differently by economists and by psychologists, and that it may refer to self-described propensity measures (e.g., personality measures), to objective behavioral tendencies in tasks specifically design to assess risk (e.g., the Balloon Analogue Risk Task) or to reports of habitual behaviors that would be categorized as risky (e.g., smoking). Presenting findings using 39 different risk-taking measures, they found evidence for a stable risk trait based on propensity and habit measures, but not behavioral tasks.

In the present study we further examine the construct of risk using data from a relatively large cohort of young adults who completed several common measures of risk-taking. In our study, we obtained several measures of risky behavior, including self-reports of perception of risk and propensity to engage in risky behaviors and performance on behavioral tasks purported to measure risk. We included three self-report questionnaires assessing perception of and propensity to take risks. These included the Survey of Consumer Finances Investment Risk Question (SCF IRQ; Aizcorbe et al., 2003), a single self-report question asking how much an individual is willing to risk financially, the Domain-Specific Risk-Taking Scale (DOSPERT; Blais & Weber et al. 2006), a multi-dimensional questionnaire assessing likelihood to engage in several other domain-specific risky activities, such “revealing a friend’s secret to someone else”, “driving a car without a seatbelt”, etc., and the Probability Choice Questionnaire (PCQ; Madden et al., 2010), which is a shorter measure that assesses self-reported preferences for smaller certain rewards over probabilistic larger rewards. Additionally, we included two behavioral tasks, one assessing choices between certain and probabilistic monetary rewards and one a designated risk-taking task. The Probability Discounting Task (PDT; Richards et al. 1997) is a behavioral task measuring actual choices between certain and uncertain (larger) rewards in a series of dichotomous choices (i.e., probability discounting behavior). The behavioral PDT and self-report PCQ assess the same underlying construct (probability discounting), but one involves behavioral choices and the other self-reported choices. The second behavioral task was the Balloon Analogue Risk Task (BART; Lejuez et al., 2002), which is a task designed to measures willingness to take monetary risk at the expense of a possible loss via inflating a simulated balloon on a computer screen. Across trials, participants may gain points with increasing risk of exploding the balloon and losing these points.

To examine the latent structure of risk preferences, we conducted a principal components analysis (PCA) to evaluate the relationship among the SCF IRQ, DOSPERT, PDT, PCQ and the BART to reveal the factor structure underlying the risk construct, and to determine how risk taking differs across domains. We used an exploratory approach (PCA) because relatively few studies have previously investigated the latent structure of risk preferences. Confirmatory factor analysis would have been more appropriate if we had specific *a priori* prediction about the nature of the structure (cf. MacKillop et al., 2016). In addition, we used PCA to examine the latent structure of all variance, not just shared variance, because we expected that risk preferences would be multi-faceted in nature. Furthermore, sex differences in risk taking behavior, across multiple domains, have been seen and replicated for decades (Fatkin et al., 1985; Powell et al., 1997; Pawlowski et al., 2008; Charness et al., 2012). Thus, we also analyzed males and females within our sample separately to account for these differences and to determine the extent to which they may influence the underlying latent structure of risk taking behaviors. Finally, a unique feature of the study was an intentional emphasis on enrolling young adults with limited involvement with drugs of abuse. Persistent substance use has been shown to lead to greater risk taking (Nasrallah et al., 2009). Therefore, using participants with non-problem drug use reduced the likelihood of either residual or long-term effects.

## 2. Methods

### 2.1 Participants

Healthy men and women (N=1058) aged 18-31 were recruited at two sites (Athens, GA and Chicago, IL) through online and printed advertisements. Online screening identified individuals who were fluent in English, had completed up to high school education, had taken no psychiatric medications in the last year, and reported no current psychiatric treatment. During the in-person visit we verified alcohol sobriety via breathalyzer (Alco-sensor III or IV; Intoximeters, St. Louis, MO) and lack of recent drug use via urine drug screen (ToxCup, Branan Medical Co. Irvine, CA and iCup, Alere North America, LLC, Orlando, FL), Participants also completed the Alcohol Use Disorder Identification Test (AUDIT) (Babor et al. 2001) and Drug Use Disorder Identification Test (DUDIT) (Berman et al. 2005), and were only included if they scored 11 or below to exclude problem drug users. The study was approved by the Institutional Review Boards of the University of Chicago and the University of Georgia, and all participants provided informed consent.

### 2.2 Procedures

Participants attended a single experimental session during which they completed self-report and behavioral measures. Participants were instructed to abstain from alcohol and drugs other than their usual amounts of caffeine and nicotine for 24 hours before the visit. Individuals with positive drug tests were excluded. The measures reported here were part of a larger battery of tasks described elsewhere (MacKillop et al. 2016). The tasks were presented in counterbalanced order, with two five-minute breaks during the 4-hour session. The present analysis consists of both self-report and behavioral indices of risk-taking (listed below). After completion of the study, participants were debriefed and compensated for their time. Participants were either paid $40 or received research participation credits, and also had a one in six chance of receiving an outcome from one of the other assessments (Kirby et al. 1999).

### 2.3 Self-report Measures

**Survey of Consumer Finances Investment Risk Question** (SCF IRQ) (Aizcorbe et al., 2003) The SCF IRQ measures financial risk-taking behavior. The single multiple-choice question asks, “which of the statements below comes closest to the amount of financial risk that you are willing to take when you save or make investments?” Possible responses included: (1) Substantial financial risks expecting to earn substantial returns, (2) Above-average financial risks expecting to earn above-average returns, (3) Average financial risks expecting to earn average returns, (4) No financial risks. This question is included in a survey sponsored by the Federal Reserve Board in cooperation with the U.S. Department of the Treasury (Grable et al. 1999).

**Domain-Specific Risk-Taking Scale** (DOSPERT) (Blais & Weber, 2006)

The DOSPERT measures risk attitudes in six commonly encountered content risk domains (ethical, gambling, health/safety, investing, recreational, and social). It is a 48-item questionnaire that assesses likelihood to engage in domain-specific risky activities, as well as perceptions of the magnitude of the risks. Sample items include “Having an affair with a married man/woman”, “Investing 10% of your annual income in a new business venture” and “Driving a car without a seatbelt”. A 7-point rating scale ranging from 1 (Extremely Unlikely) to 7 (Extremely Likely) was used to assess likelihood to engage in the stated risky behavior. Item ratings were added across all items of a given subscale to obtain subscale scores. Higher scores indicate greater risk taking in the domain of the subscale. The risk-perception assessment used the same set of items but instead included a 7-point rating scale ranging from 1 (Not at all) to 7 (Extremely Risky). Item ratings were added across all items of a given subscale to obtain subscale scores, with higher scores suggesting perceptions of greater risk in the domain of the subscale.

**Probability Choice Questionnaire** (PCQ) (Madden et al., 2010) The PCQ measures probability discounting behavior. Participants are instructed to answer 30 questions by circling their preference between two outcomes, a smaller amount of money delivered “for sure” and a probabilistic larger amount, in no particular order. For example, one item asks participants “Would you rather have $20 for sure or a 1-in-10 chance (10%) of winning $80”. Each probability reflects predetermined discounting functions, which permit inferring a value for the parameter *h*.

### 2.4 Behavioral tasks

**Probability Discounting Task** (Richards et al. 1997)

The PDT measures the relative value of certain vs probable consequences. A computerized procedure was used to present choices in which participants repeatedly chose between $100 with a probability (1.0, 0.9, 0.75, 0.5 and 0.25) and a smaller, certain amount. Indifference points, at which two options are perceived as equal in value to an individual, are used to plot discount curves. The curve represents the rate of the probability discounting and is best characterized by a hyperbolic model. The hyperbolic discount functions for probability discounting are calculated as follows:

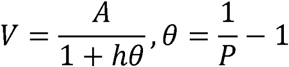

The V represents the subjective value (the certain smaller amount of money), the A represents the larger amount money ($100). The P represents the probability of receiving the money, the 0 stands for odds against receiving the money, and the *h* represents the rate of discounting as a function of decreasing probability. Lower *h* values represent a less rapid rate of discounting based on increasing odds against, reflecting riskier options.

**Balloon Analogue Risk Task** (BART) (Lejuez et al., 2002)

The BART is a validated behavioral measure of risk taking (Hunt et al., 2005, Lejuez et al., 2007). Participants view a balloon on a screen, which can be increased in size with a key press. Each key press increases the balloon size and increases a counter on the screen, with points redeemable for money. However, as the balloon increases in size the probability that it will explode also increases, at which time all accumulated points are lost. The subject can make an alternative response to stop pumping before the balloon explodes and redeem the points. Thus, this task provides a measure of willingness to take risk, at the expense of a possible loss. The adjusted average number of pumps on unexploded balloons is the indicator of risk.

### 2.5 Data Analysis

Initially, for all measures, distributions were examined, and log transformations were performed when necessary to normalize skewed data. Sex differences across all measures were determined. Then, internal reliability and correlations between measures and their subscales were evaluated. Finally, the latent structure of the measures of risk taking was examined using principal components analysis (PCA). To identify related latent factors, the PCA used an oblique rotation (direct oblimin, δ = 0), permitting correlated components. Two criteria were used to determine the appropriate number of components to retain: eigenvalues > 1, and scree plot discontinuity. Significant loadings were defined as >|.30| on the pattern matrix. All data analysis was performed in SPSS (v24).

## 3. Results

### 3.1 Participant Characteristics

The participants were mostly young men and women of European-ancestry with about 2 years of college education (Table 1).

**Table 1.**
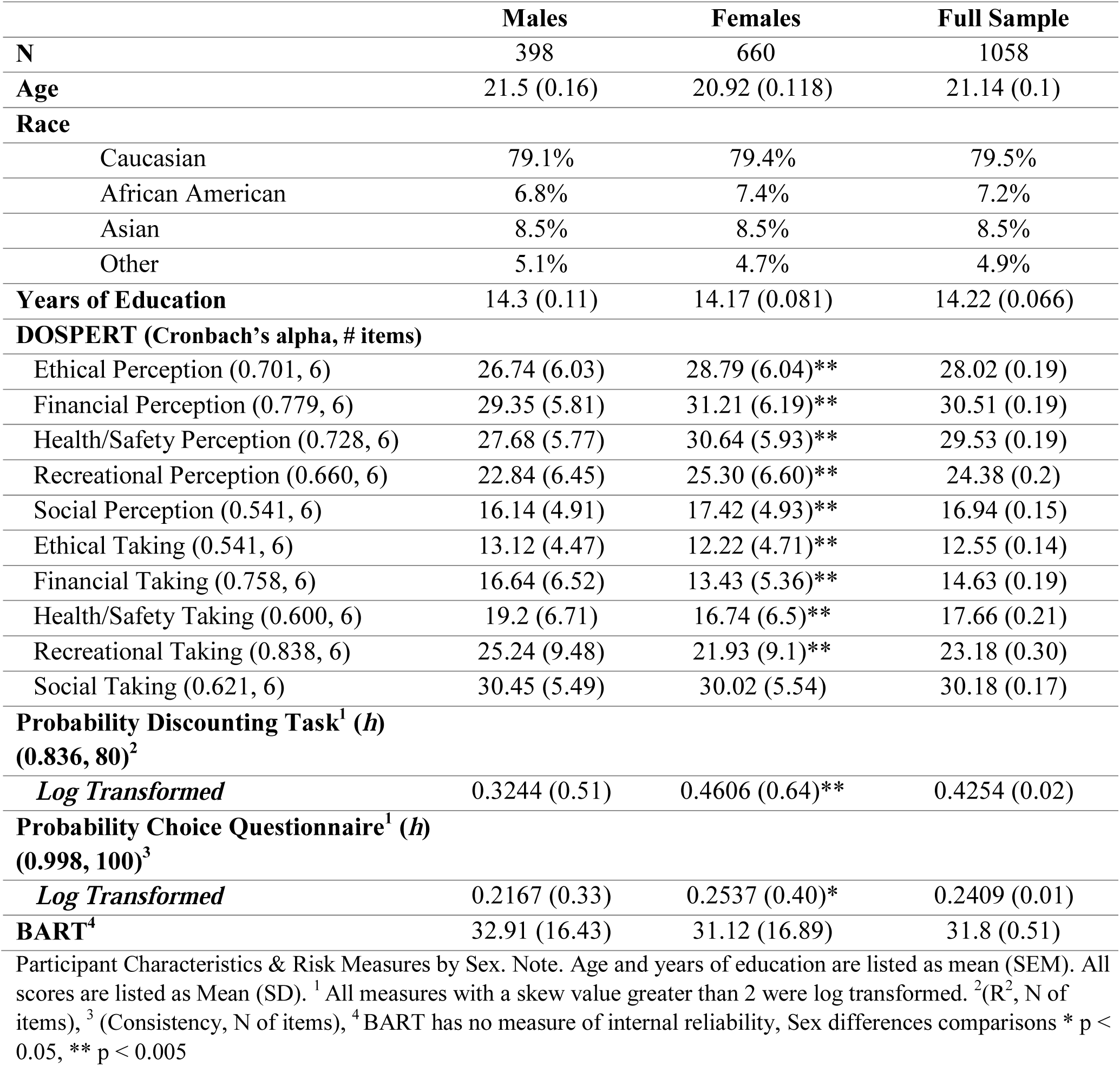
Participant Characteristics & Risk Measures by Sex. Note. Age and years of education are listed as mean (SEM). All scores are listed as Mean (SD). ^1^ All measures with a skew value greater than 2 were log transformed. ^2^(R^2^, N of items), ^3^ (Consistency, N of items), ^4^ BART has no measure of internal reliability, Sex differences comparisons * p < 0.05, ** p < 0.005

### 3.2 Self-Report & Behavioral Measures

Subjects’ mean and standard deviations for each of the outcome measures (DOSPERT subscale scores, *h* values, and mean unexploded balloons adjusted) are listed in Table 1. These values are within the normative range reported for the DOSPERT (Blais & Weber, 2006), the probability discounting task (Richards et al. 1997), the probability choice questionnaire (Madden et al., 2010) and the BART (Lejuez et al., 2002). Women and men differed on all DOSPERT subscales except social risk-taking behavior, on both the probability task and questionnaire, but not on the BART. Females perceived all domains of the DOSPERT risk-taking subscales, except social, to be significantly riskier than males, but all scored significantly lower on likelihood to take the risks in all domains in comparison to males (Table 1). Additionally, because of these observed sex differences, we conducted the primary analyses in separate phases, first for the entire sample, then adjusted for sex and finally separately for females and males.

### 3.3 Preliminary Analyses

Before conducting the principal components analysis, we examined both the internal reliability of the DOSPERT (Table 1) and the correlations between measure subscales. The *h* values of both probability discounting and probability choice measures were significantly correlated (*r*s > 0.34, *p* < 0.005) and inherently reflect the same content, and therefore were combined into a single measure by taking the arithmetic mean. The DOSPERT perceived risk and likelihood scores for each subscale were also correlated (*r*s > 0.37, *p* < 0.005) and therefore multiplied, yielding a composite of ‘risk orientation’, representing the weighting of risk attribution by likelihood of engaging in it.

### 3.4 Full Sample Analysis

Principal components analysis was conducted on the full sample utilizing an oblique rotation. The analysis yielded three components (Table 2). The first component accounted for 27.3%, the second 14.3% and the third 12.7%, for a total of 54.3% of the variance. Component 1 included DOSPERT ethical, health/safety, recreational and social risk and was thus labeled “*Lifestyle Risk Sensitivity*.” Component 2 consisted of three variables, the combined probability discounting measures, the inverse of DOSPERT financial risk and the SCFIRQ, and was thus labeled “*Financial Risk Sensitivity*.” Component 3 included the BART alone and was thus labeled “*Behavioral Risk Sensitivity*.”

**Table 2.** Principal Components Analysis. All values over 0.3 are in bold. Note. component loadings taken from the pattern matrix

| Measure Outcome | Full Sample Loadings |  |  | Sex-Adjusted PCA Loadings |  |  | Male PCA Loadings |  |  | Female PCA Loadings |  |  |
| --- | --- | --- | --- | --- | --- | --- | --- | --- | --- | --- | --- | --- |
|  | Component 1: Lifestyle Risk Sensitivity | Component 2: Financial Risk Sensitivity | Component 3: Behavioral Risk Sensitivity | Component 1: Lifestyle Risk Sensitivity | Component 2: Financial Risk Sensitivity | Component 3: BART Risk Sensitivity | Component 1: Lifestyle Risk Sensitivity | Component 2: Financial Risk Sensitivity | Component 3: Behavioral Risk Sensitivity | Component 1: Lifestyle Risk Sensitivity | Component 2: Financial Risk Sensitivity | Component 3: BART Risk Sensitivity |
| Combined Probability Discounting | 0.02 | **0.546** | -0.149 | 0.022 | **0.56** | -0.208 | 0.267 | **0.704** | 0.054 | -0.090 | **0.478** | **0.325** |
| DOSP ERT Ethical Risk | **0.557** | -0.181 | -0.043 | **0.558** | -0.181 | -0.02 | **0.319** | -0.239 | **-0.409** | **0.639** | -0.025 | -0.066 |
| DOSP ERT Financial Risk | 0.277 | **-0.643** | -0.168 | 0.298 | **-0.607** | -0.203 | **0.530** | **-0.445** | 0.074 | 0.296 | **-0.633** | 0.212 |
| DOSP ERT Health & Safety Risk | **0.745** | 0.003 | 0.198 | **0.758** | 0.042 | 0.206 | **0.595** | -0.146 | -0.067 | **0.800** | 0.244 | -0.231 |
| DOSP ERT Recreational Risk | **0.662** | -0.126 | -0.034 | **0.672** | -0.094 | -0.044 | **0.778** | 0.022 | -0.114 | **0.581** | -0.199 | 0.034 |
| DOSP ERT Social Risk | **0.645** | 0.209 | -0.071 | **0.628** | 0.156 | -0.076 | **0.660** | 0.259 | 0.082 | **0.539** | -0.026 | 0.266 |
| SCFIR Q | 0.075 | **0.814** | 0.069 | 0.075 | **0.812** | 0.093 | -0.249 | **0.707** | -0.065 | 0.175 | **0.824** | 0.009 |
| BART | 0.044 | -0.009 | **0.962** | 0.043 | -0.012 | **0.941** | 0.051 | -0.131 | **0.946** | 0.028 | -0.022 | **-0.872** |

### 3.5 Sex Adjusted Analysis

Principal components analysis was then conducted on the full sample using sex-adjusted measures of each risk measure. Sex adjusted measures were created by regressing each risk measure with sex and saving the standardized residuals. The analysis yielded three components (Table 2). The first component accounted for 27.2%, the second 14.0% and the third 12.7%, for a total of 53.9% of the variance. Component 1 included DOSPERT ethical, health/safety, recreational and social risk and was thus labeled “*Lifestyle Risk Sensitivity*.” Component 2 consisted of three variables, the probability discounting measure, the DOSPERT financial risk and the SCFIRQ, and was thus labeled “*Financial Risk Sensitivity*.” Component 3 included the BART alone and was thus labeled “*Behavioral Risk Sensitivity*.”

### 3.6 Sex Specific Analysis

Principal components analysis was conducted on males and females separately. For both males and females, the analysis yielded three components (Table 4). For females, the first component accounted for 24.9% the second 15.4% and the third 12.9%, for a total of 53.3% of the variance. Component 1 included DOSPERT ethical, health/safety, recreational and social risk and was thus labeled “*Lifestyle Risk Sensitivity*.” Component 2 consisted of the combined probability discounting measures, the inverse of DOSPERT financial risk, and the SCFIRQ, and because these all involved finances this component was labeled “*Financial Risk Sensitivity*.” Component 3 included the BART alone and was thus labeled “*Behavioral Risk Sensitivity*”

For males, the first component accounted for 28.8% the second 14.1% and the third 12.7%, for a total of 55.5% of the variance. Component 1 included DOSPERT financial, health/safety, recreational, ethical and social risk and was thus labeled “*Lifestyle Risk Sensitivity*.” Component 2 consisted of two variables, the combined probability discounting measures and the SCFIRQ, and because these both involved money this component was labeled “*Financial Risk Sensitivity*.” Component 3 included BART and the DOSPERT ethical risk subscale was thus labeled “*Behavioral Risk Sensitivity*”.

## 4. Discussion & Conclusion

This principal component analysis of several indices of risk-taking revealed a latent structure of three distinct components of risk-taking behavior. Based on their content, we labeled these: “*Lifestyle Risk Sensitivity*”, which included ethical, recreational, health and safety, and social risk, “*Financial Risk Sensitivity”*, which included the two measures of monetary risk as measured by probability discounting and “*Behavioral Risk Sensitivity”*, which consisted only of the BART. Males and females differed on most of the measures: Males reported to be more likely to engage in risky behaviors, and females reported a higher perception of risk. A sex-adjusted principal components analysis revealed an identical three component structure to the full sample analysis. Similar components were extracted for both males and females analyzed separately, except that the DOSPERT ethical subscale loaded onto the *Behavioral Risk Sensitivity* component in males, but not females. Overall, our findings support the existence of three similar underlying components of risk-taking behavior measures: *Lifestyle, Financial and Behavioral Risk Sensitivity*.

These findings support the current view that risk-taking is multidimensional. Although risk-taking was previously considered a unitary personality trait (i.e., risk-taking or risk averse) a growing body of evidence suggests that risk-taking consists of different components, or domains (Blais & Weber, 2001; 2002; 2006). Our findings support this idea, indicating that risky lifestyle behaviors (health, safety etc.) may not be related to risky financial behaviors, and that the BART assesses a separate, unrelated dimension.

Furthermore, our findings add to our understanding of sex differences within risk taking behaviors. Typically, males are more inclined to take risks across a variety of domains, including financial to lifestyle, and females are more risk averse (Fatkin et al., 1985; Powell et al., 1997; Pawlowski et al., 2008; Charness et al., 2012). This was also seen in our results. Females perceived all domains of the DOSPERT risk subscales, except social, to be significantly riskier than males, and they scored significantly lower on likelihood to take the risks in all domains compared to males. Thus, females were more risk averse and perceived risks as riskier. Interestingly the sex-adjusted PCA indicated nearly the same three components as our full sample unadjusted PCA, suggesting similar underlying factor structure in men and women. When PCA’s were conducted separately for men and women we found one minor difference in component loadings—the DOSPERT ethical variable loaded onto “*Behavioral Risk Sensitivity”* rather than “*Lifestyle Risk Sensitivity”* in males but not in females. Whether this difference is specific to this sample or reflects a general sex difference is not known.

Similar to more recent work evaluating the Balloon Analogue Risk Task, in our study, behavior on the BART was not related to other indices of risk. During the development of the BART, Lejuez et al (2002) showed that it was correlated with several real-life risk-taking behaviors including addictive, health, and safety risk behaviors. Since then there have been numerous reports that the BART differentiates populations thought to be risk-takers from non-risk-takers (e.g., smokers vs nonsmokers, individuals high on self-reported impulsivity or psychopathy, jailed inmates, cocaine users; Lejuez et al. 2003, Hunt et al. 2005, Lejuez et al. 2007, Swogger et al. 2010, Tull et al. 2009). However, these relationships have not always been consistently present and some studies have even reported opposite relationships (Courtney et al. 2012; Ryan et al. 2013). Furthermore, prior work has also showed that neither the BART nor the Iowa Gambling Task were correlated with self-report measures of impulsivity (BIS-11, I-7 Impulsivity, MPQ Constraint) or sensation seeking (I-7 Venturesomeness; Reynolds et al., 2006). Reynolds et al. concluded that “self-report and behavioral tasks probably measure different constructs, and … even among the behavioral measures, different tasks measure different, perhaps unrelated, components of impulsive behavior” (Reynolds et al., 2006, p. 305–306). The present study is one of the first to investigate the relations among survey and behavioral measures of risk-taking using a latent variable approach in a well-powered sample of adolescents. The findings supported the discrepancy between the BART, a behavioral task, and self-report tasks by revealing the unique latent component BART loaded onto by itself.

Within the financial risk category, our findings with probability discounting raised a methodological issue: We measured probability discounting using both a behavioral task and a self-report questionnaire, and the results were highly correlated. In this case, the distinction between ‘behavioral’ measure and ‘self-report’ measure is not completely clear because the content of the choices using the two methods was similar (i.e., certain smaller amount of money vs larger probabilistic amount of money). Yet, the high correlation between the measures suggests that the participants’ behavior was driven by the content rather than the form of the measure. In our study, both probability discounting measures were also correlated with the self-report measure of financial risk-taking, suggesting that there is a general financial risk-taking underlying construct that is dissociable from non-financial risk. Our findings diverge slightly from the findings by Frey et al (2017), who used many more measures (39 measures, compared to our 7) to derive the factor structure of different indices of risk. They found a weak correlation between propensity (self-report) and behavioral measures of risk, but they did find an overall general risk factor that was related to frequency of engaging in real-life risky behaviors like smoking. Of course, the problem with examining smoking and other risk substance use is that it includes processes that may in fact be a cause of risk phenotypes.

Our study extended knowledge about risky behaviors in several ways. First, we identified three distinct constructs reflecting apparently unrelated forms of risk-taking behavior: lifestyle, financial and behavioral risk sensitivity. Notably, the financial construct was comprised of both self-report measures and behavioral tasks, providing good support for a true underlying construct. Second, we ascertained these constructs in participants who were relatively homogeneous in terms of age, absence of psychiatric symptomatology or addictive behaviors, thereby minimized the possible confounds that these variables might contribute to the data. At the same time, however, the relative homogeneity of our sample does raise a question about whether these same constructs would exist in more mixed populations. In addition, although we used an extensive battery of measures, not every measure of risk preference was represented. Future work utilizing more, and different measures of risk-taking behavior, both self-report and behavioral, such as the Risk Perception Scale (Benthin et al., 1993), in addition to the DOSPERT, and the Wheel of Fortune Task (Ernst, M. et al., 2004), in more heterogeneous samples, such as individuals of different ethnicities and with an illicit drug use history, will help to identify the true biobehavioral constructs underlying these behaviors.

## Acknowledgements

We thank Drs. Jessica Weafer and Joshua Gray for their contributions to the data collection and guidance throughout.

**Funding** This work was partially supported by National Institutes of Health grant R01 DA032015 (de Wit, Palmer, MacKillop) and the Peter Boris Chair in Addictions Research (MacKillop).

## Supplementary Material

**Supplementary Table 1.**

| Measure | Combined Probability Discounting | DOSPERT Ethical | DOSPERT Financial | DOSPERT Health/safety | DOSPERT Recreational | DOSPERT Social | SCFIRQ | BART |
| --- | --- | --- | --- | --- | --- | --- | --- | --- |
| Combined Probability Discounting | **1.000** |  |  |  |  |  |  |  |
| DOSPERT Ethical | **-0.132\*\*** | **1.000** |  |  |  |  |  |  |
| DOSPERT Financial | **-0.131\*\*** | **0.284\*\*** | **1.000** |  |  |  |  |  |
| DOSPERT Health/safety | **-0.106\*** | **0.362\*\*** | **0.194\*\*** | **1.000** |  |  |  |  |
| DOSPERT Recreational | **-0.117\*\*** | **0.196\*\*** | **0.326\*\*** | **0.354\*\*** | **1.000** |  |  |  |
| DOSPERT Social | -0.003 | **0.151\*\*** | **0.148\*\*** | **0.184\*\*** | **0.264\*\*** | **1.000** |  |  |
| SCFIRQ | **0.162\*\*** | **-0.125\*\*** | **-0.397\*\*** | **-0.092\*\*** | **-0.142\*\*** | -0.048 | **1.000** |  |
| BART | -0.005 | -0.050 | -0.038 | 0.039 | -0.030 | -0.013 | -0.007 | **1.000** |

Full sample correlation matrix. Note. Zero-order Pearson's correlations among measures and subscales of risk taking, \* p<.05, \*\* p<.005

**Supplementary Table 2.**

| Measure | Combined Probability Discounting | DOSPERT Ethical | DOSPERT Financial | DOSPERT Health/safety | DOSPERT Recreational | DOSPERT Social | SCFIRQ | BART |
| --- | --- | --- | --- | --- | --- | --- | --- | --- |
| Combined Probability Discounting | **1.000** | **-0.144\*\*** | **-0.138\*\*** | **-0.103\*** | **-0.166\*\*** | -0.006 | **0.137\*\*** | -0.016 |
| DOSPERT Ethical | **-0.108\*** | **1.000** | **0.272\*\*** | **0.393\*\*** | **0.147\*\*** | **0.165\*\*** | -0.039 | -0.008 |
| DOSPERT Financial | -0.086 | **0.309\*\*** | **1.000** | **0.105\*** | **0.272\*\*** | **0.168\*\*** | **-0.296\*\*** | -0.037 |
| DOSPERT Health/safety | **-0.100\*** | **0.314\*\*** | **0.293\*\*** | **1.000** | **0.313\*\*** | **0.212\*\*** | 0.052 | 0.076 |
| DOSPERT Recreational | -0.014 | **0.269\*\*** | **0.387\*\*** | **0.409\*\*** | **1.000** | **0.256\*\*** | **-0.079\*** | 0.013 |
| DOSPERT Social | -0.023 | **0.135\*\*** | **0.187\*\*** | **0.161\*\*** | **0.300\*\*** | **1.000** | -0.073 | -0.006 |
| SCFIRQ | **0.176\*\*** | **-0.246\*\*** | **-0.467\*\*** | **-0.267\*\*** | **-0.207\*\*** | -0.069 | **1.000** | -0.039 |
| BART | 0.032 | **-0.121\*\*** | -0.07 | -0.03 | **-0.107\*** | -0.011 | 0.067 | **1.000** |

Correlation matrix by sex. Note. Zero-order Pearson's correlations among measures and subscales of risk taking, Males below middle diagonal, Females above middle diagonal, \* p<.05, \*\* p<.005

**Supplementary Table 3.**

| **Component** | *Lifestyle Risk Sensitivity* | *Financial Risk Sensitivity* | *Behavioral Risk Sensitivity* |
| --- | --- | --- | --- |
| *Lifestyle Risk Sensitivity* | **1.000** | -0.01 | -0.078 |
| *Financial Risk Sensitivity* | -0.204** | **1.000** | -0.21** |
| *Behavioral Risk Sensitivity* | -0.077 | -0.022 | **1.000** |

**B.**
| **Component** | *Lifestyle Risk Sensitivity* | *Financial Risk Sensitivity* | *Behavioral Risk Sensitivity* |
| --- | --- | --- | --- |
| *Lifestyle Risk Sensitivity* | **1.000** | 0.004 | 0.010 |
| *Financial Risk Sensitivity* | -0.161* | **1.000** | -0.181** |
| *Behavioral Risk Sensitivity* | -0.113* | 0.161** | **1.000** |

Component Correlations: Sex Specific Samples Note. Zero-order Pearson's correlations among measures and subscales of risk taking, Females upper right hand part of table, Males lower left hand part of table, \* p<.05, \*\* p<.005

**Supplementary Figure 1.**
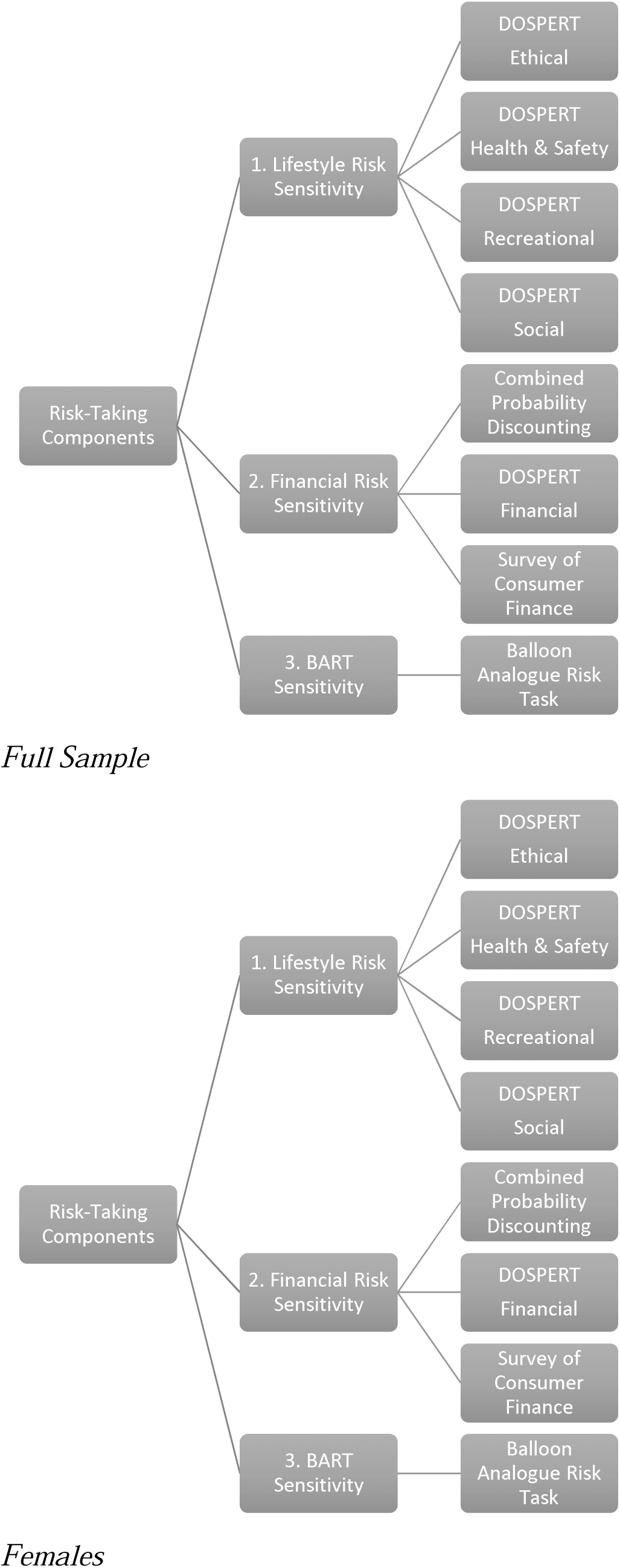
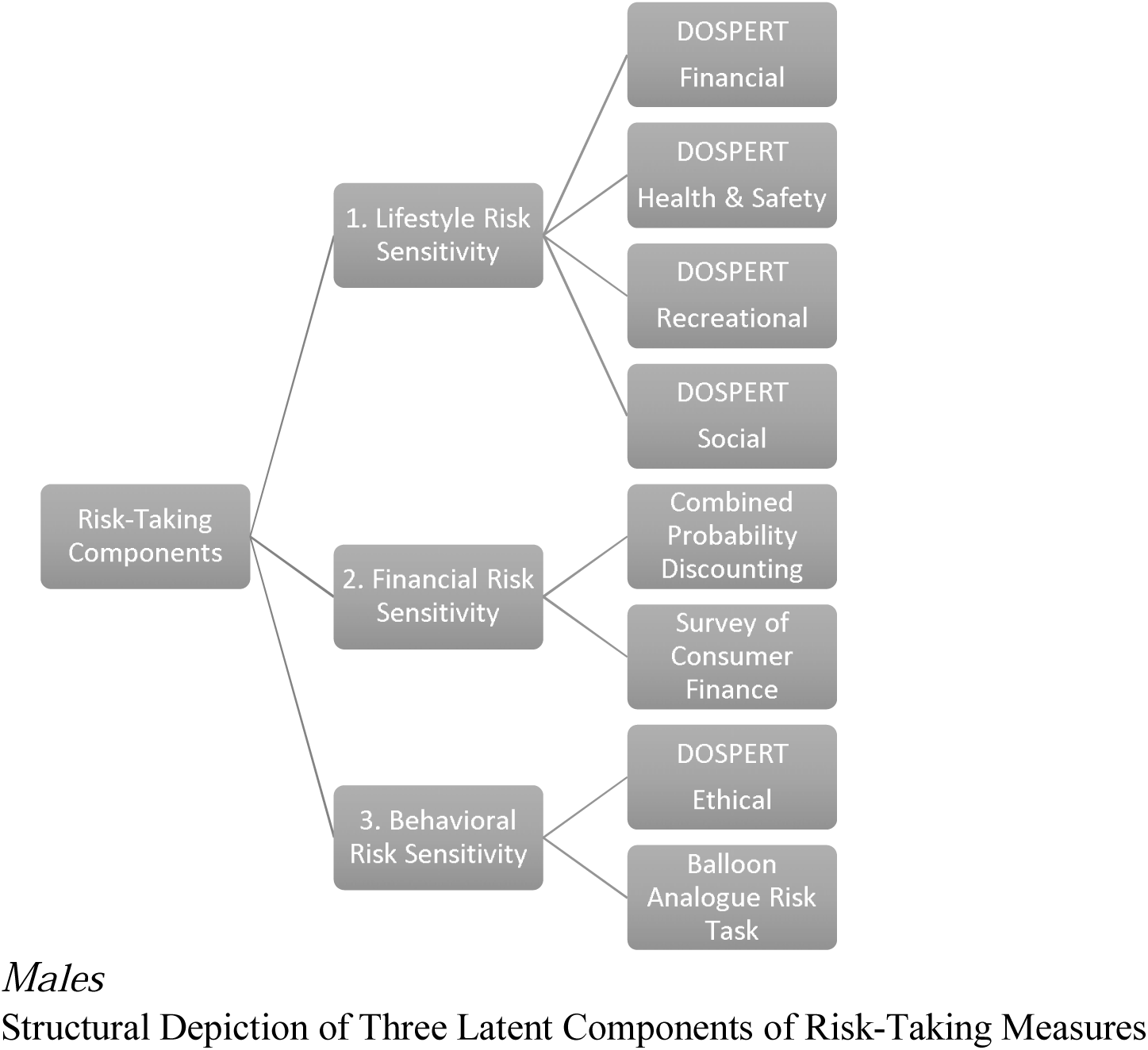

